# The packaging signal of *Xanthomonas* integrative filamentous phages

**DOI:** 10.1101/2023.12.30.573655

**Authors:** Ting-Yu Yeh, Patrick J. Feehley, Michael C. Feehley, Vivian Y. Ooi, Pei-Chen Wu, Frederick Hsieh, Serena S. Chiu, Yung-Ching Su, Maxwell S. Lewis, Gregory P. Contreras

**Author notes:** These authors contributed equally (Patrick J. Feehley, Michael C. Feehley, Vivian Y. Ooi). Corresponding author: Ting-Yu Yeh. This study is dedicated to Dr. Pien-Chien (P.C.) Huang (1931-2020), a professor of the Johns Hopkins University and Academician of Academia Sinica, for his deep devotion to the students and their education for more than 50 years.

## Abstract

Unlike Ff, the packaging signal (PS) and the mechanism of integrative filamentous phage assembly remain largely unknown. Here we revived two *Inoviridae* prophage sequences, ϕLf2 and ϕLf-UK, as infectious virions lysogenize black rot pathogen *Xanthomonas campestris pv. campestris*. ϕLf2 and ϕLf-UK genomes consist of 6,363 and 6,062 nucleotides and share 85.8% and 98.7% identity with ϕLf, respectively. To explore their assembly, we first identified 20-26– nucleotide long PS sequences of 10 *Xanthomonas* phages. These PS consist of a DNA hairpin with the consensus GGX(A/-)CCG(C/T)G sequence in the stem and C/T nucleotides in the loop, both of which are conserved and essential for PS activity. In contrast to Ff, the 5’ to 3‘ orientation of PS sequence is not conserved or critical for their competence. This is the first report to offer insights into the structure and function of integrative phage PS, revealing the diversity of filamentous phage encapsidation.

## Introduction

*Xanthomonas*, a group of approximately 27 Gram-negative bacterial species, are the major phytopathogens with significant economic importance (Ryan et al., 2011). *Xanthomonas* species cause bacterial spots and blights on the leaves, stems, and fruits of nearly 400 plant species. We and others have reported several *Xanthomonas* filamentous phages of *Inoviridae* family, including *Xanthomonas oryzae* (Xf, Xf2, ϕXo, and Xf109), *Xanthomonas citri* (Cf, XacF1 and Cf2), *Xanthomonas campestris pv. campestris* (Xcc, ϕLf), and *Xanthomonas campestris pv. vesicatoria* (ϕXv and XaF13) (Kuo et al., 1969; Dai et al., 1980; Kamiunten and Wakimoto, 1980; Tseng et al., 1990; Lin et al., 1994; Ahmad et al., 2014; Yeh, 2017; Solis-Sanchez, et al., 2020; Yeh, 2020). Unlike *Escherichia coli* Ff phages (f1, fd, M13, etc.) that replicate as episomes, all *Xanthomonas* filamentous phages are often lysogenized by the integration of their genomes into the host bacterial chromosomes, where they replicate as prophages (Kuo et al., 1987a; Kuo et al., 1987b; Dai et al., 1987; Dai, et al., 1988; Bai, 1989; Fu et al., 1992; Lin et al., 1994; Kamiunten, 1995; Ahmad et al., 2014; Yeh, 2017; Yeh, 2020). Cf (Cflt and Cf16) is the first “integrative” filamentous phage reported (Kuo et al., 1987a; Kuo et al., 1987b; Dai et al., 1987). Since then, integrative filamentous phages have been identified in variety of human, animal and plant pathogens, and they play crucial roles in disease pathogenesis and virulence of their host bacteria (review by Mai-Prochnowet al., 2015; Hay and Lithgow, 2019). The best example is the filamentous phage CTXϕ whose genome encodes the cholera toxin during *Vibrio cholerae* infection, causing life-threatening diarrhea (Waldor and Mekalanos, 1996).

*Xanthomonas campestris pv. campestris* (Xcc) causes black rot disease in brassica crops worldwide, which is considered the most destructive disease of crucifers (Williams PH. 1980). ϕLf is the first filamentous phage identified in Xcc, and its complete nucleotide (nt) sequence is available in NCBI GenBank (MN263053) (Tseng et al., 1990; Wen and Tseng, 1994; Lin et al., 1996; Wen and Tseng, 1996; Liu, et al., 1997). Here we report the characterization and complete sequences of two new integrative filamentous phages, ϕLf-UK and ϕLf2, revived from the Xcc strain ATCC 33913 and 8004, respectively.

The structures of coliphage (fd, If1, M13) and other filamentous phages (Xf, Pf1, Pf3, Ike) have been well characterized by different tools (X-ray diffraction, NMR, cryo-electron microscopy, etc.) (Day, et al., 1988; Connors, et al., 2023; Jia and Xiang, 2023). Ff phage assembly starts from the packaging signal (PS), which is a DNA stem loop. PS binds to pVII and pIX (the “round end”) at the bacterial inner membrane and translocate through Zot ATPase pI/pXI. Major coat protein pVIII is added, and the phage protrudes through the pIV channel at the outer membrane. Other minor coat proteins pIII and pVI (the “pointy tip”) are added to terminate this process (Hay and Lithgow, 2019; Connors, et al., 2023). Even though it has been over 50 years since the discovery of the first *Xanthomonas* filamentous phage Xf (Kuo et al., 1969), how *Xanthomonas* or other integrative filamentous phages assemble, mature, and release their viral particles remains poorly understood.

Genomic evidence shows that the pIV homologue is missing from most integrative filamentous phages in Gram-negative bacteria, such as in the *Vibrio cholerae* phage CTXϕ, the *Ralstonia solanacearum* phage (ϕRSM1 and ϕRSS1), the *Stenotrophomonas maltophilia* phages (ϕSMA6, ϕSMA7, ϕSMA9, ϕSHP1, ϕSHP2) and all the *Xanthomonas* phages (Cf1c, XacF1, Xf109, Xf409, Cf2, ϕLf, XaF13, ϕXv2 and ϕLf-UK and ϕLf2 in this study). In addition, many filamentous phages (e.g. *Xanthomonas* or *Stenotrophomonas*) lack the Ff pIX homologue (as well as pVII), which binds to the PS to initiate the phage assembly cascade. These observations prompted us to explore how these phages are encapsidated. To answer this question, we first teased out the PS motif from the *ori* sequence reported in previous works (Lin et al., 1996; Lin and Tseng, 1996). We identified the eight other integrative *Xanthomonas* filamentous phage PS, which share sequence and structural similarity. Many molecular and structural properties of these PS are different from the well characterized Ff PS, suggesting that filamentous phage assembly/egress may be different across the varieties of filamentous phages.

## Materials and Methods

### Bacteria and plasmids

To generate the host bacteria Xcc-TcR derived from Xcc ATCC 33913 (ATCC, Manassas, VA, USA), 12,151 base pair (bp) ϕLf-homologous sequence from the chromosome DNA (AE008922.1; 2435696-24478046) was replaced by the tetracycline resistant gene from the *Pfo*I/*Eco*RI fragment of pRGD-TcR (Addgene, #74110) (Ledermann et al, 2016) through double crossover, then the bacteria was selected by tetracycline as described (Lin, et al., 2001). pORIPS plasmid, which contains the ϕLf-UK *ori* and PS sequence (245-365, MH206184; identical to ϕLf fragment by Lin and Tseng, 1996) was synthesized and subcloned into pOK12 plasmid containing the kanamycin resistance gene (Vieira and Messing, 1991). The ORF1 gene with synonymous substitutions at the PS (T340A/C341G) and the *ori* (nt 277-318; GA**A** GG**A** CA**A A**G**A** GCA GC**A T**T**A** GC**A T**T**A** GG**A T**T**A** GT**T** CAT TA**T**, mutations in bold) of ϕLf-UK was amplified by PCR and subcloned into a broad-host-range expression, chloramphenicol resistance vector pHP11 (Reece and Phillips, 1995) as pGII plasmids. Plasmids were purified by Plasmid Maxi Kit (Qiagen) according to the manufacture’s protocol. Site directed mutagenesis of pORIPS and pGII was performed by PCR using the QuikChange II Site-directed mutagenesis kit (Agilent).

### Phage revival and characterization

Prophage sequences of ϕLf-UK (Xcc ATCC 33913; AE008922.1; 2435698-2441774) and ϕLf2 (Xcc 8004; CP000050.1, 2537488-2543837) were synthesized and cloned in the *Bam*HI site of the pUC57-Kan vector (GenScript, Piscataway, NJ) and propagated in *E. coli*. Plasmid DNAs were purified and digested with *Bam*HI. The prophage fragments were recovered from the 1% (w/v) agarose gel, re-ligated with T4 DNA ligase, and then electroporated into Xcc-TcR (Wang and Tseng 1992). Phage propagation, purification of phage particles, phage and replicative form (RF) DNA, and host chromosomal DNA were described accordingly (Yeh, 2020). Phage morphology, bacterial growth after phage infection, and phage yield were characterized as previously (Tseng et al., 1990; Lin et al., 1994; Yeh, 2020).

Phage integration was analyzed by PCR using oligonucleotide primers (IDT) as described (Yeh, 2017; Yeh 2020). Primers sequences for *att*B, *att*P, *att*L, and *att*R are Xcc-TcR (#1: 5’-CTTACGCATCTGTGCGGTATTTC-3’, #4: 5’-GCACCAAGATTACAGCCAGCC-3’), ϕLf-UK (#2: 5’-GAATTCACGCAGAACGAGCGG-3’, #3: 5’-GCCTTCGGGCGTGATGAGGTG-3’), ϕLf2 (#5: 5’-GATGAGGTGGCCACCCTGGAAC-3’, #6: 5’-GCAGCTGCATCGCCGGCTGAG-3’).

Computer-aided assembly of the nucleotide sequence data was performed with SerialCloner 2-5 program. Database searches assigning the open reading frames (ORFs) and gene functions were analyzed with the FASTA, BLASTN, BLASTP, and Clustal Omega programs. The complete genome sequences of ϕLf2 and ϕLf-UK phage were deposited in DDBJ/ENA/GenBank under the accession number MH218848 and MH206184, respectively.

PS DNA structure was predicted by RNAFold (Gruber et al., 2008). Prediction of transmembrane helices was carried out by DeepTMHMM (Hallgren et al, 2022). Signal peptides were predicted by SignalP 5.0 (Almagro et al., 2019). The phylogenetic tree was generated by FigTree software (version 1.4.2, Andrew Rambaut, University of Edinburgh, UK) after rooting with an outgroup protein sequence of the *Stenotrophomonas maltophilia* (WP_240796989) as described previously (Rambaut, 2021; Yeh and Contreras, 2021; Yeh et al. 2022).

### Ori activity

1 ng of pGII plasmid was co-transformed with pORIPS plasmid (1 ng) into Xcc-TcR by electroporation. 50% and 50% of the transformed bacteria were plated in a kanamycin (50 µg/ml) and chloramphenicol (20 µg/ml) selective plate, respectively, and grown at 28 °C for 16 h. The *ori* activity was determined by colony numbers of pORIPS transformants in the kanamycin plates normalized with the chloramphenicol selection.

### Detection of kanamycin-resistant transducing particles

To generate kanamycin-resistant transducing particles (TPs), Xcc-TcR cells were transformed with pORIPS plasmid and plated on a LB plate with tetracycline (12 µg/ml) and kanamycin (50 µg/ml) at 28°C for 18 h. Three colonies were picked, and each colony was incubated in one milliliter of culture at 28°C for 18 h. pORIPS transformants were superinfected with ϕLf-UK phage (multiplicity of infection = 20). Infected bacterial cultures were centrifuged, and the supernatant was filtered through a 0.45-µm-pore-size filter. The titers of pORIPS TPs in the filtrate were measured by infecting Xcc-TcR as described previously (Lin et al 1996).

### Determination of phage concentration using SYBR Green quantitative PCR (qPCR) method

The copy numbers of pORIPS phage particles released in medium was measured by qPCR based on Peng et al.’s method (Peng et al., 2018). The ϕLf-UK phage ssDNA and pORIPS plasmid (concentration 1 μg/μL) were diluted into a serial concentration of 1 fg/μL–10^7^ fg/μL, which was used to prepare the standard curve for titers of genome copies of ϕLf-UK phage and pORIPS. The following equation was applied to calculate the standard DNA concentration from fg/μL to genome copies per microliter (gc)/μL:

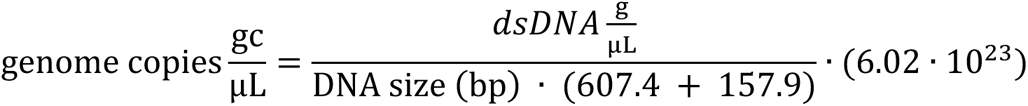

Phages from the filtrates described above were precipitated with 5% polyethylene glycol 6000 and 0.5 M NaCl on ice for 4 hours and centrifuged at 13,5000 *g* for 15 min at 100 °C. The phage pellets were resuspended in TE buffer (10 mM Tris, pH 8.0, 1 mM EDTA) and diluted into ten-serial, ten-fold concentrations. Residual DNA not from intact phage particles was removed before DNA extraction from phages for qPCR (Peng et al., 2018). Phage samples were pre-treated with 2.5 μL of DNase I (2,000 units/mL, NEB), 2.5 mM MgCl_2_ and 0.5 mM CaCl_2_ in 200 μL of each diluted concentration of phage samples at 37 °C for 10 min. DNase I-treated samples were heat-denatured at 100 °C for 15 min. Each concentration of denatured phage samples was subjected to qPCR.

PowerUp SYBR Green Master Mix (Cat. # A25742) was purchased from ThermoFisher Scientific. qPCR primers were designed by PrimerQuest Tool (IDT). Primers (IDT) for ϕLf-UK phage were forward 5’-CGCTGTCGGCAATTCTCTAT-3’, reverse 5’-ATGGTGCTACCCGCATTAC –3’. Primers for pORIPS TP DNAs were forward 5’-TGCGCCAGAGTTGTTTCT-3’, and reverse 5’-GATGGTCGGAAGAGGCATAAA-3’. A 10 µL PCR reaction mixture included 5 µL 2 x SYBR Green Master Mix, 1 µL 0.5 mM primers, 2 µL H2O and 2 µL denatured phages samples. PCR conditions were 50 °C for 2 min, 95 °C 2 min, 40 cycles of 95 °C for 15 s, and 58 °C for 1 min, followed by the melting curve setting of 1 cycle of 95 °C for 15 s, 58 °C for 1 min, and 95 °C for 15 s. The copy numbers of pORIPS TP DNAs were normalized with ϕLf-UK phage released in the medium. Each condition was triplicated, and at least three colonies were tested for each pORIPS mutant.

## Results

### Revival of two filamentous Xcc phages, ϕLf-UK and ϕLf2

We first transformed the prophage sequences of ϕLf-UK (Xcc ATCC 33913; AE008922.1; 2435698-2441774) and ϕLf2 (Xcc 8004; CP000050.1, 2537488-2543837) into Xcc-TcR host to revive and amplify the phage. After CsCl centrifugation purification, ϕLf-UK and ϕLf2 phage particles were filamentous in shape and measured approximately 1010 (±150) × 8 nm (ϕLf-UK) and 1070 (±185) × 8 nm (ϕLf2) (n=100, Figure 1A and 1B). SDS-polyacrylamide gel electrophoresis of the purified virions showed a major single band of approximate *M*_r_ 4,000, like Cf2 reported previously (Fig. 1C). Similar to ϕLf but in contrast to Cf2 and Xf109, these phages multiplied without retarding host cell growth rate at 16 hours post-infection (p.i.) (Yeh, 2017; Yeh, 2020). By infection of 10^8^ ml^-1^ bacteria at a multiplicity of infection of 20, viral particles of ϕLf-UK and ϕLf2 were produced at the titer of 2.1-3.2×10^11^ and 3.5-5.2×10^11^ PFU ml^-1^ at 16 hours p.i.

**Figure 1.**
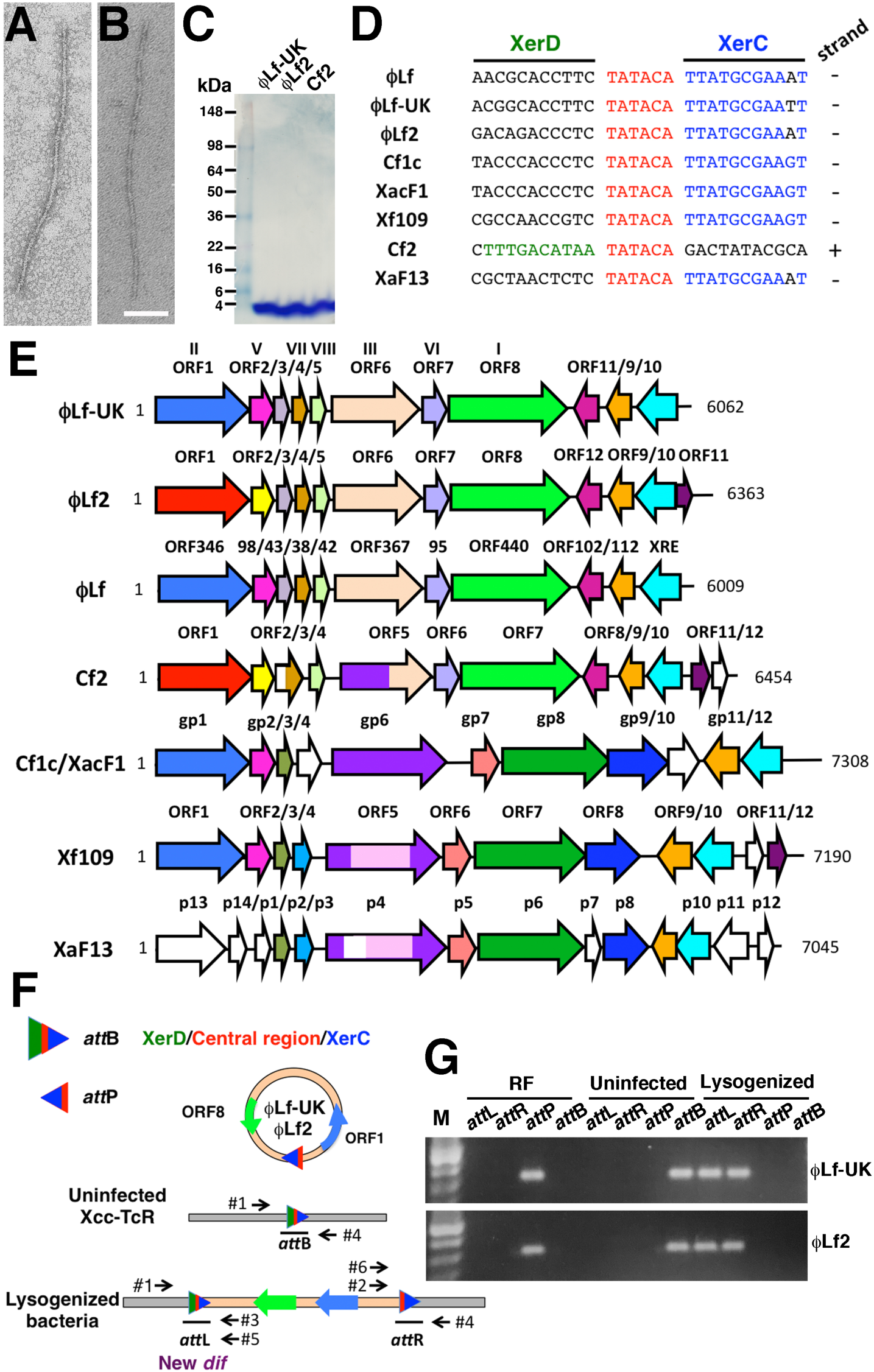
Characterization of ϕLf-UK and ϕLf2 phages and genomes. (A and B) Electron micrograph of ϕLf-UK (A) and ϕLf2 (B) phage negatively stained with 2% (w/v) phosphotungstic acid. Bar=100 nm. (C) Purified phages were subjected to 4-20% SDS-polyacrylamide gel electrophoresis (SDS-PAGE) followed by staining with Coomassie blue. (D) The *att*P sequences *Xanthomonas* filamentous phages. XerD-binding arm (green), central region (red), XerC-binding arm (blue), and their locations on the positive or negative strand of phage RF DNAs are also indicated. (E) Genomic organization of *Xanthomonas* filamentous phages. Arrows oriented in the direction of transcription represent ORFs. (F and G) Site-specific integration of ϕLf-UK and ϕLf2 phage DNA into the host chromosome. (F) PCR strategy for *att*P sequence in RF, the *att*B sequence in uninfected cells, and *att*L and *att*R sequences in the lysogenized host. PCR primers are used for Xcc-TcR (#1 and #4), ϕLf-UK (#2 and #3), and ϕLf2 (#5 and 6). ORF1 and ORF8 are labeled (not to scale) for the phage genome orientation. The XerD-binding arm (green), central region (red) and XerC-binding arm (blue) are indicated. (G) PCR fragments of ϕLf-UK (upper) and ϕLf2 (lower) as (F) in a 1% (w/v) agarose gel in Tris-borate-EDTA buffer with the DNA marker (M).

The phage lysogeny was confirmed by PCR using primers (Figure 1F and 1G), which flank 350 base-pair fragments of Xcc-TcR chromosomal DNAs containing *att*L and *att*R (Figure 1G). Phage production in ϕLf-UK– and ϕLf2-lysogenized cell cultures was 1.1-2.3×10^4^ and 3.5-5.4×10^4^ PFU ml^-1^, respectively. Taken together, infectious progeny phage particles of ϕLf-UK and ϕLf2 can be revived and lysogenize the host bacteria *de novo*.

### Genome organization of ϕLf-UK and ϕLf2 phage

ϕLf2 and ϕLf-UK phage genomes consist of 6,363 and 6,062 nucleotides (nts), which sequences (98% and 70% query coverage for ϕLf-UK and ϕLf2) share 98.7% and 85.8% identity with ϕLf (MN263053, Wen and Tseng, 1994; Lin et al., 1996; Wen and Tseng, 1996; Liu, et al., 1997), respectively. The major coat protein B (ORF5) and the *dif* sequences (5’-TATACATTATGCGAA-3’) of ϕLf-UK and ϕLf2 are identical to ϕLf, suggesting that these three Xcc phages are closely related. The genome feature of ϕLf-UK and ϕLf2 phage is organized in the order gII-gV-gVII-gVIII-gIII-gVI-gI (ORF1-2-4-5-6-7-8) on the positive strand (Figure 1E, see later) (Liu, et al., 1997). Their minus strands contain a helix-turn-helix XRE-family DNA binding protein, a DUF3653 domain protein, and a putative chloride channel protein, which all are conserved in ϕLf and Cf2 (Peng, 2002, Yeh, 2020).

The ϕLf-UK genome contains 11 predicted open reading frames (ORFs) (Figure 1E, Table 1). The major difference between ϕLf-UK and ϕLf is located at ϕLf-UK ORF9 and ϕLf ORF112 (nt 263-601, X70328.1), which AAs and nucleotides share 62% and 66% identity, respectively. In addition, ϕLf-UK (ORF6) encodes minor coat protein pIII homologue with extra 19 amino acids in the glycine-rich linker region compared to ϕLf (Wen and Teng, 1996).

**Table 1.** Predicted ORFs in ϕLf2 and ϕLf-UK phage genome.

| Coding Sequence | Strand | Position 5'-3' | Amino acids | Molecular mass (Mr.) | Amino acid sequence identity/similarity to homologues (no. of amino acid identical; % identity) | E-value | Accession number |
| --- | --- | --- | --- | --- | --- | --- | --- |
| **ϕLf2** |  |  |  |  |  |  |  |
| **ORF1** | + | 1-1089 | 362 | 40,344.69 | ϕSHP2 phage *RstA* (256; 72) | 0.0 | YP_004539131 |
| **ORF2** | + | 1086-1382 | 98 | 10,824.29 | ϕSHP2 phage *RstB* (65; 68) | 6e-46 | AED02384 |
| **ORF3** | + | 1414-1545 | 43 | 4,709.56 | ϕLf phage ORF43 (42; 98) | 9e-29 | CAA49797 |
| **ORF4** | + | 1521-1637 | 38 | 4,044.69 | ϕLf phage ORF38, minor coat protein gVII (38; 100) | 2e-28 | CAA49798 |
| **ORF5** | + | 1665-1793 | 42 | 4,114.87 | ϕLf phage ORF42, major coat protein B, gVIII (42; 100) | 1e-27 | CAA49799 |
| **ORF6** | + | 1869-2939 | 356 | 35,401.74 | ϕLf phage ORF367, phage absorption protein, gIII (135; 73) | 8e-97 | AAB87092 |
| **ORF7** | + | 2950-3270 | 106 | 11,931.37 | ϕLf phage ORF95, minor coat protein B, gVI (91; 93) | 5e-67 | AAB88260 |
| **ORF8** | + | 3272-4594 | 440 | 47,923.51 | ϕLf phage ORF440, phage zonula occludens toxin, gI (412; 94%) | 0.0 | AAB88261 |
| **ORF12** | - | 4617-4922 | 102 | 11,548.37 | Cf2 ORF8 chloride channel protein Eric (91; 88) | 2e-66 | QBJ05107 |
| **ORF9** | - | 4989-5315 | 108 | 11,922.81 | ϕLf phage ORF112 (44; 60) | 2e-30 | CAA49794 |
| **ORF10** | - | 5316-5813 | 166 | 18,116.11 | DUF3693 family, Cf1c cp8, helix-turn-helix XRE-family DNA binding protein, (130; 92) | 2e-78 | NP_536679 |
| **ORF11** | + | 5812-6171 | 119 | 12,655.49 | Cf2 ORF11 protein (53; 76) | 5e-24 | QBJ05110 |
| **ϕLf-UK** |  |  |  |  |  |  |  |
| **ORF1** | + | 1-1041 | 346 | 39,172.72 | ϕLf phage ORF346, replication initiation protein, gII (346; 100) | 0.0 | U38235 |
| **ORF2** | + | 1038-1334 | 98 | 10,990.52 | ϕLf phage ORF98, ssDNA-binding protein, phage gV protein (98; 100) | 0.0 | X70331 |
| **ORF3** | + | 1366-1497 | 43 | 4,683.48 | ϕLf phage ORF43 (43; 100) | 0.0 | X70331 |
| **ORF4** | + | 1473-1589 | 38 | 4,044.69 | ϕLf phage ORF38, minor coat protein gVII (38; 100) | 0.0 | X70331 |
| **ORF5** | + | 1617-1745 | 42 | 4,114.87 | ϕLf phage ORF42, major coat protein B, gVIII (42; 00) | 0.0 | X70331 |
| **ORF6** | + | 1773-2933 | 386 | 38,366.74 | φLf ORF367, minor coat protein A, gIII (367; 95) | 0.0 | U10884 |
| **ORF7** | + | 2968-3264 | 98 | 11,014.44 | φLf phage ORF95, minor coat protein B, gVI (79, 99) | 5e-56 | AF018286 |
| **ORF8** | + | 3266-4588 | 440 | 48,136.69 | φLf phage ORF440, phage zonular occludens toxin, gI (440; 100) | 0.0 | AF018286 |
| **ORF11** | - | 4611-4916 | 102 | 11,548.37 | Cf2 ORF8 chloride channel protein EriC (91; 88) | 2e-66 | QBJ05107 |
| **ORF9** | - | 4983-5309 | 108 | 11,921.82 | φLf phage ORF112 (42; 62) | 5e-31 | X70328 |
| **ORF10** | - | 5299-5844 | 181 | 19,631.81 | Cf1cp8, helix-turn-helix XRE-family DNA binding protein (82; 64) | 4e-47 | NP_536679 |

The ϕLf2 genome consists of 12 predicted ORFs (Figure 1E, Table 1). ϕLf2 replication proteins *Rst*A/*Rst*B share no homology to ϕLf gII/gV (Figure 1E and Table 1). The nucleotide sequences of ϕLf2 ORF1 and ORF2 are 74% identical to *Stenotrophomonas* phages ϕSHP2 or ϕSMA7 (NC015586 and NC021569; Liu, et al, 2012; Petrova et al., 2014). The ϕLf2 ORF1 and ORF2 protein share 72% and 68% amino acid homology to ϕSHP2 *Rst*A and *Rst*B, but only 33% and 40% to Cf2 ORF1 and ORF2. ϕLf2 has an additional ORF11 protein absent in ϕLf-UK or ϕLf, while this homologue is present in Cf2, Xf109, and *Stenotrophomonas* phage ϕSMA7. This suggests that the ϕLf2 genome may have evolved from multiple *Xanthomonas* and *Stenotrophomonas* phage ancestors, likely via genetic recombination (Yeh, 2020).

### Are gVII and gIX present in integrative filamentous phages?

Based on early work and recent cryoEM studies, pVII and pIX form the “round end” of M13 and f1 phage particles (Endemann and Model 1995; Connors, et al., 2023; Liu and Xiang, 2023). It is believed that pVII and pIX are the first proteins secreted during assembly of the phage particle (Lopez and Webster, 1983). They are transmembrane proteins that bind to PS DNA prior to translocation of phage DNA at pI/pXI machinery (Russel and Model, 1989; Mai-Prochnow, et al., 2015; Hay and Lithgow, 2019). Previously it has been proposed that ϕLf ORF43 and ORF38 are Ff gVII and gIX homologues (Wen and Tseng 1994, Liu et al., 1997). After revisiting the sequence alignment and AlphaFold structural prediction of these two proteins, we found that ϕLf ORF38, Cf2 ORF3, ϕLf-UK and ϕLf2 ORF4 encode pVII rather than previously proposed pIX (Figure S1A and S1B). Based on DeepTMHMM prediction, there is a transmembrane domain between their amino acid 7 to 29, which was also seen with AlphaFold (Figure S1B). ϕLf ORF38 homologues’ N-termini usually contain aspartic acid or glutamic acid residues together, and the last amino acid in their C-termini is a conserved arginine, which are all features conserved in gVII of coliphages or *Salmonella* phage Ike (Figure S1A). Phylogenetic analysis also supports that ϕLf ORF38 is related to gVII. (Figure S2)

Although ϕLf ORF43 was originally proposed as gVII, ϕLf ORF43 and the ORF3 of Cf2, ϕLf-UK and ϕLf2 do not contain any transmembrane domain or share any sequence homology with gVII or gIX. This protein appears to have a putative lipoprotein signal peptide at its N-terminus, and the cleavage site is predicted to be located between amino acid 17 and 18 (Figure S1C and S1D). This result suggests that this protein may be transported by the Sec translocon and cleaved by signal peptidase II Lsp, although its function requires further study.

Taken together, we conclude that ϕLf, ϕLf-UK, ϕLf2, and Cf2 have gVII homologues but not gIX. Moreover, gIV, gVII and gIX are absent in Cf1c, XacF1, Xf109, Xf409, and XaF13 genomes (Figure 1E). gVII and gIX are also not present in other integrative filamentous phages. Our data suggest the “leading/round end” essential for viral assembly could be very different between Ff and integrative filamentous phages.

### Xanthomonas filamentous phage PS

Loss of well-known Ff phage assembly proteins (gIV, gIX, gVII) in these genomes leads us to investigate how *Xanthomonas* or other integrative filamentous phages generate their virial particles. Since Ff gVII and gIX interact with the PS DNA, we first explored PS properties of *Xanthomonas* phage. It has been shown that a ϕLf fragment (identical to nt 245-365 of ϕLf-UK) can support phage DNA replication and miniphage production, suggesting that it contains both a replication origin (*ori*) and a PS for encapsidation (Lin and Tseng, 1996). However, the exact PS sequence of ϕLf or any other integrative filamentous phages has not been identified in detail so far. Therefore, we chose ϕLf-UK as our model virus in this study.

Based on the thermodynamic ensemble of the centroid structure using RNAFold software (Gruber et al, 2008), this fragment consists of 4 stem-loop (SL) like structures (SL1: 245-272; SL2: 273-306, SL3: 307-331, SL4: 332-357) (Figure 2A). The conserved “recognition sequence” 5’-CTTG-3’ and putative nicking site G304 for the superfamily I Rep protein (gII) is located at SL2 (Lin and Tseng, 1996. The putative “key binding” sequence of *ori* is located at SL3 (Koonin and Ilyina, 1993). SL3 also contains a GNA trinucleotide loop (GCGNAGC), which is extraordinarily stable. (Yoshizawa et al., 1997)

**Figure 2.**
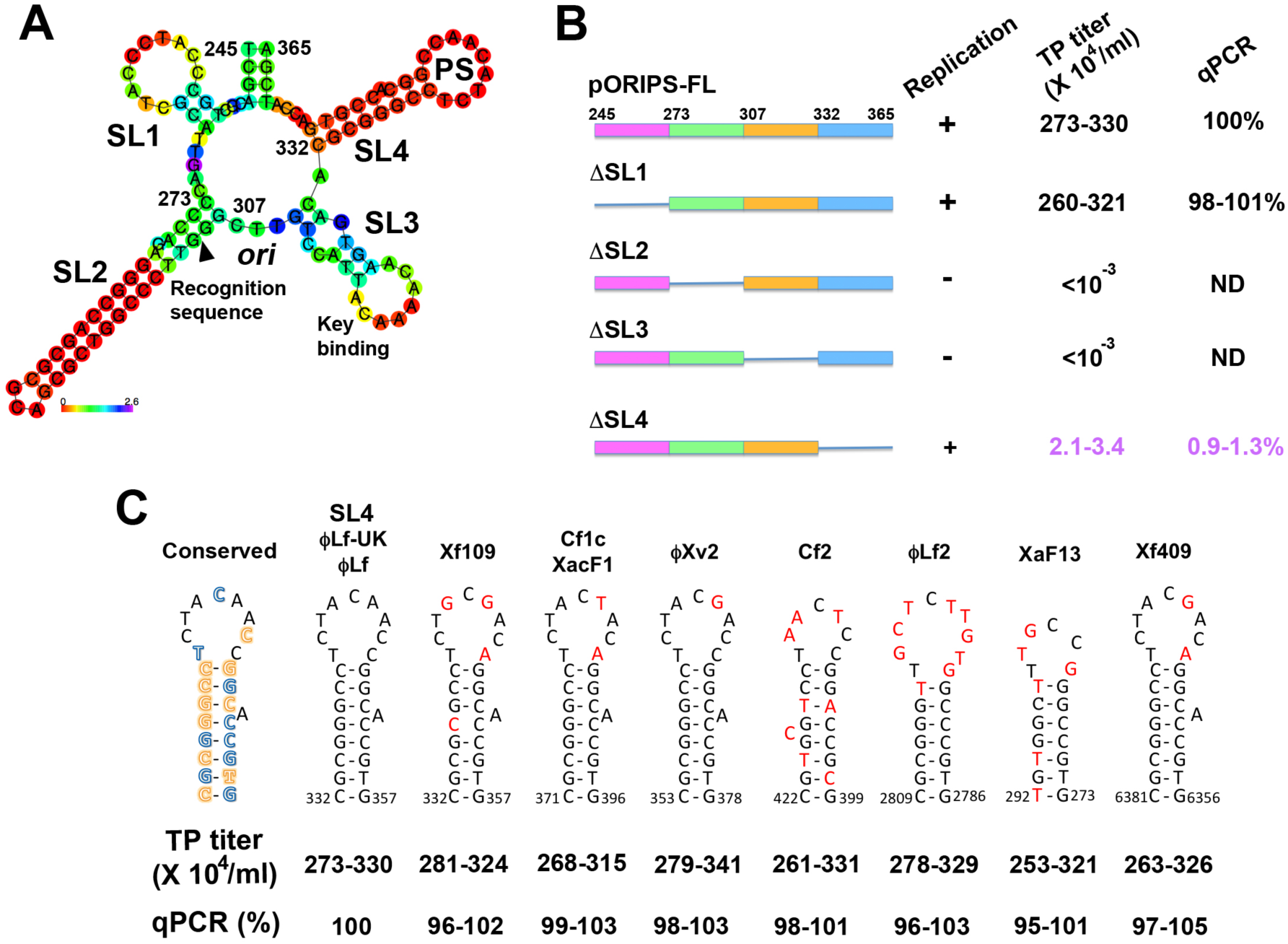
Characterization of *Xanthomonas* phage PS. (A) Thermodynamic ensemble of the centroid structure of ϕLf-UK nt 245-365 using RNAFold. The positional entropy for each position was present. (B) Mapping of *ori* and PS sequences in (A). *ori* activity was assayed by colony formation of pORIPS transformants under kanamycin selection. PS activity was determined by TP titers and the DNA copy numbers of pORIPS TPs released in media. The copy number of pORIPS-FL TP DNA normalized by the copy number of ϕLf-UK helper phage was taken to be 100%. ND for “not detected”. (C) Schematic representation of the secondary structure of *Xanthomonas* filamentous phages PS, whose position was according to NCBI sequences (ϕLf-UK: MH206184; ϕLf2: MH218848; Xf109: KX181651; Cf2: MK512531; XacF1/Cf1c: AB910602; ϕXv2: MH206183; XaF13: MN335248; Xf409: KY853667. Please note that complement Xf409 sequence was deposited.). Nucleotide variations are shown in red. Completely and >80% conserved nucleotides of 10 *Xanthomonas* phages PS were labeled in blue and orange, respectively. PS activity was assayed as (B). The results of (B) and (C) were from three independent experiments, 3 colonies for each experiment.

To further map the ϕLf-UK *ori* and PS, we first constructed the pORIPS plasmid by subcloning ϕLf-UK nt 245-365 (full-length, FL) or serial SL deletions (ΔSL1 to ΔSL4) in a pOK12 vector, which contain the kanamycin resistance gene but cannot replicate as an episome in *Xanthomonas* (Figure 2B; Yeh, 2020). The *ori* replication efficiency was determined by co-transforming pORIPS derivatives with the chloramphenicol-resistant pGII plasmid which encode ϕLf-UK ORF1 (gII) protein into XCC-TcR. The phage PS and *ori* in pGII were mutated to avoid competition with pORIPS. The transformants were selected by chloramphenicol and kanamycin.

Colony numbers in kanamycin selection plates were not affected in ΔSL1 and ΔSL4 (97-103 transformant/ng pORIPS plasmid), but almost abolished in ΔSL2 and ΔSL3 (< 10^-2^/ng) (Figure 2B, three experiments). Our result indicates that ϕLf-UK and ϕLf *ori* are located in SL2 and SL3, including the putative recognition and key binding sequences of gII.

To test phage production ability, XCC-TcR was transformed with pORIPS derivatives and then superinfected with ϕLf-UK phage. The titers of pORIPS TPs in the culture medium were analyzed by infecting XCC-TcR, followed by screening for kanamycin resistance. pORIPS-FL TP titer was approximately 2.7 to 3.3 ×10^6^/ml, comparable to previously reported ϕLf pOR1 (2.6 ×10^6^ /ml) or pT2 (0.8-2.3 ×10^6^ /ml) TPs (Lin et al., 1996; Lin and Tseng, 1996). We did not find colony formation of *ori* mutants ι1SL2 and ι1SL3, consistent with their replication deficiency. ι1SL1 had normal TP titer (2.6 to 3.2 ×10^6^/ml), but ι1SL4 TP titer drastically reduced to 2.1-3.4 ×10^4^/ml, about 1% of pORIPS-FL. This result suggests that SL4 is required for either TP generation (packaging) or its infectivity (Figure 2B). To rule out SL4 role in phage infectivity, we also analyzed the DNA copy number of pORIPS TPs released in the culture medium using qPCR. Our data confirmed that ι1SL2 and ι1SL3 DNA were not detected in the culture medium, and ι1SL4 TP DNA was around 100-fold (0.9-1.3%) less compared to pORIPS-FL. Taken together, we concluded that the ϕLf-UK/ϕLf SL4 is PS crucial for efficient encapsidation but not replication.

ϕLf-UK PS is a DNA hairpin (nt 332-357) with a GC-rich stem and dyad symmetry (Figure 2A and 2C). We next examined whether the PS sequences or structures are conserved among 8 other *Xanthomona*s filamentous phages (ϕLf, ϕLf-UK, ϕLf2, Cf1c, Cf2, XacF1, Xf109, Xf409, ϕXv2, XaF13) with their complete genome sequences. The homologous sequences of 20-26 nucleotides SLs can be identified in the gII of Xf109, Cf1c, XacF1, Xf409, and ϕXv2 (Figure 2C). Interestingly, the “chiral” DNA SL sequences with the opposite 5’ to 3’ direction can be also identified in Cf2, ϕLf2, and XaF13. These SLs are also located at the coding region of replication initiation protein *Rst*A of Cf2 ORF1 and replication protein A of XaF13 ORF13, respectively, except one of ϕLf2 at ORF6 (gIII homologue). To verify whether these SLs act as PS, we replaced pORIPS SL4 with them and measured the TP titers and DNAs in culture media. All *Xanthomona*s phage SLs, but not the negative control antisense PS (PS-) (Figure 3A), can fully support TP reproduction as FL (Figure 2C), indicating that they are PS and functionally interchangeable.

**Figure 3.**
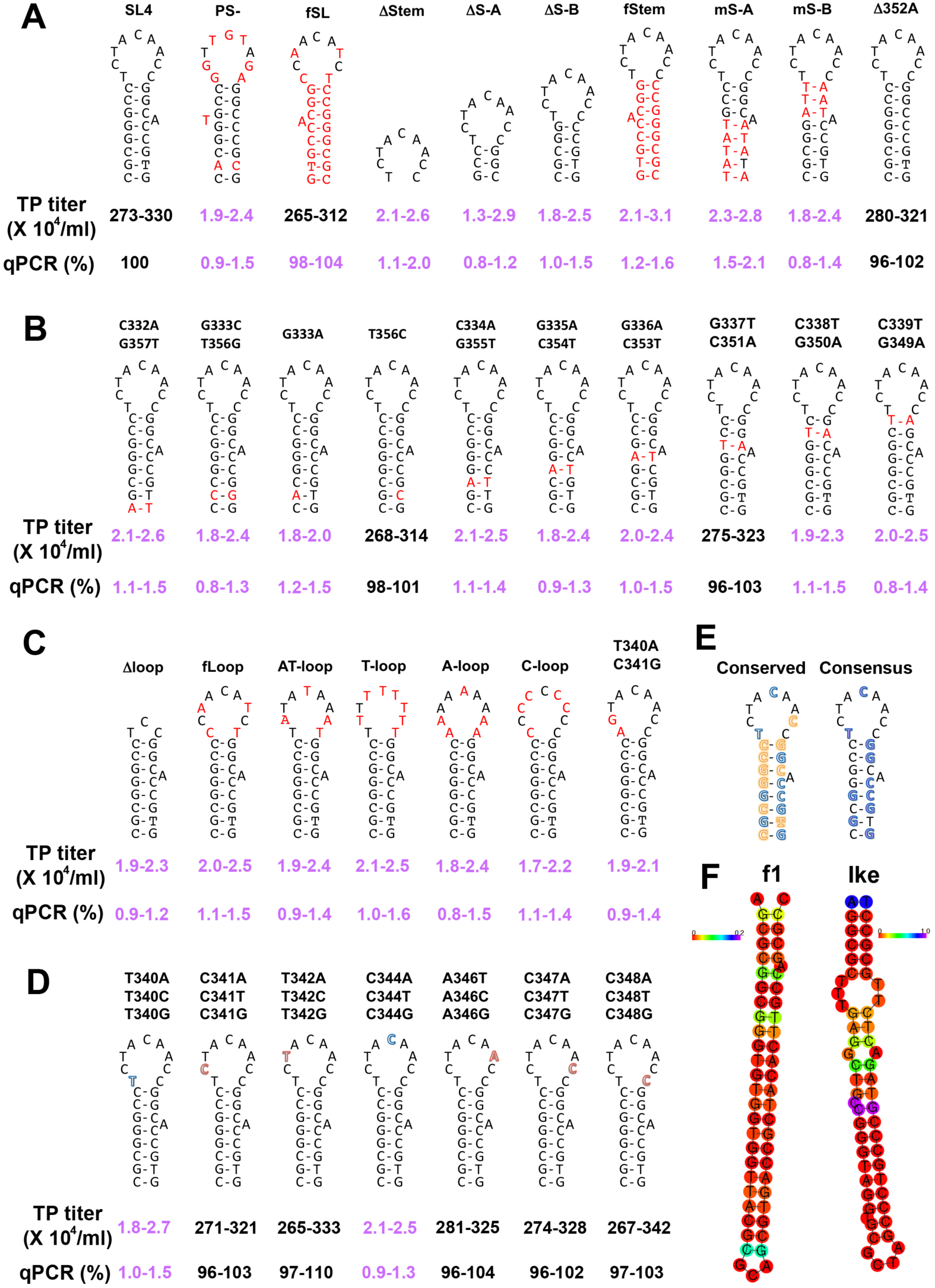
Sequence and structural requirement of ϕLf-UK PS activity. (A-D) Mutational analysis of ϕLf-UK PS. Stem (A and B) and loop (C and D) mutations were assayed for PS activity as described in Figure 2B. The results were from three independent experiments, 3 colonies for each experiment. (E) Comparison of conserved PS nucleotides (blue and orange, Figure 2C) and consensus nucleotides essential for PS activity (purple). (F) Centroid structure drawing of 54 nt f1 PS (NC_025823, nt 5511-5564) and 57 nt Ike (NC_002014, nt 6141-6197) PS encoding positional entropy based on RNAFold.

### Essential structures and sequences for Xanthomonas filamentous phage PS activity

All *Xanthomonas* phage PS contain a 7-8 base pairs GC-stem with conserved G333, G335, G350, G353, C354, G355, G357 nucleotide. Their loops have 6-10 nts with two conserved nucleotides, T340 and C344 (Figure 2C). This led us to hypothesize that these sequence and structural motifs are essential for ϕLf-UK PS function. To test this hypothesis, we performed site-directed mutational analysis of SL4 in the pORIPS-FL (Figure 3). Full or partial deletions (ΔStem, ΔS-A, ΔS-B) or replacements of C-G base pairs with A-T base pairs (mS-A, mS-B) in the stem significantly impaired PS activity (0.8-2.1%), indicating that the length of GC-rich stem plays an essential role (Figure 2A). T365C, G337T/C351A, or Δ352A did not affect PS activity, but TP titers were severely diminished in other stem mutants (G333A, C332A/G357T, G333C/T356G, C334A/G355T, G335A/C354T, G336A/C353T, C338T/G350A, C339T/G349A) (Figure 3A and 3B). These results indicate that the consensus “GGX(A/-)CCG(C/T)G” sequence of the stem is required for ϕLf-UK PS competence (Figure 3E).

Two conserved nucleotides, T340 and C344, are essential because substitutions at entire loop (AT-loop, T-loop, A-loop, C-loop) or at T340 (T340A, T340C, T340G, T340A/C341G) and C344 (C344A, C344T, C344G) abolished PS activity. Because substitutions at other loop position (C341, T342, A346, C347 and C348) had no effect, the deleterious effects were position and nucleotide specific (Figure 3C and 3D). The importance of T340 and C344 was also supported by the fact that PS-sequence still contains the consensus stem sequence (GGCACCGCG) but with T340G/C344G mutations in the loop. pORIPS TPs yield were also low when only TCC nucleotides are at the tip (Δloop, Figure 2C), suggesting that T/C nucleotides alone are not sufficient to restore PS function. Our mutagenesis data also showed these consensus nucleotides in the stem and loop are also highly conserved among *Xanthomonas* phages PS regardless of its 5’ to 3’ orientation (Figure 3E).

Dotto et al. have shown that f1 PS can only function when it is inserted in the same orientation as *ori*. (Dotto, 1981). The fact that PS at both orientations among *Xanthomonas* phages led us to examine whether the orientation of PS sequences matters. We found that encapsidation was not affected with the opposite 5’ to 3’ direction of full ϕLf-UK PS sequence (fSL) (Figure 2A). Furthermore, PS activity also remained when the sequence orientation was changed in eight other *Xanthomonas* phages PS (data not shown). Surprisingly, orientation changes only in stem or loop (fStem and fLoop) failed to restore PS activity (Figure 2A and 2C). Therefore, although *Xanthomonas* phage PS orientation is not critical for assembly, the spatial structure relationship between the stem and loop may still be important for PS functional competence.

## Discussions

It has been well established that lysogenic phages are frequently involved in bacterial virulence, including *Yersinia* phage CUSϕ-2 (Gonzalez et al, 2002), and *Vibrio* phages VGJϕ (Campos et al., 2003) and CTXϕ (Huber and Waldor, 2002). Filamentous phages play a crucial but usually underappreciated role in many other human, animal or plant diseases. Here we reported that two lysogenic phages, ϕLf-UK and ϕLf2, can infect the black rot disease pathogen Xcc. They are the second filamentous phages identified in Xcc with complete sequence information. The influence of ϕLf-UK and ϕLf2 infection on Xcc virulence remains to be further investigated. Emerging genomic information provides the opportunity to explore their evolutionary and functional diversity.

Filamentous phage PS is a *cis*-acting element of the viral genome located near the leading end of the phage particle, where it determines the orientation of DNA within the virion during encapsidation (Webster et al., 1981; Lopez and Webster, 1983). Based on RNAFold software, both f1 (54 nt) and Ike (57 nt) PS are predicted to form a long stem-loop structure (Figure 3F). Functional mappings and genetic analysis of f1 and Ike PS has suggested the required sequence and orientation necessary for its encapsidation activity (Peeters et al., 1985; Russel and Model, 1989). Our mapping indicated that SL2 and SL3 are *ori’*s, and SL4 acts as a PS. Mutation analysis further revealed the consensus nucleotides and structural requirement for *Xanthomonas* phage PS activity. To our knowledge, this is the first experimental evidence that identifies PS structural motifs (the stem and loop) involved in integrative filamentous phage assembly.

There are many differences between the Ff and *Xanthomonas* phage PS in this study. First, the 5’ to 3’ orientation of PS sequences does not affect the production yield of *Xanthomonas* filamentous phages, but the orientation is critical for Ff and Ike phages (Dotto, et al., 1981, Webster et al., 1981; Shen et al., 1979). Second, the efficient encapsidation of ϕLf-UK requires T340 and C344 residues at the PS loop, while substitution or insertion at the loop did not affect Ff PS functions (Russel and Model, 1989). Third, except for ϕLf2, the *Xanthomonas* phage PS are located at the ORF where it encodes the replication proteins (gII, *Rst*A, replication gene A protein), while for Ff or *Salmonella* Ike phage, the PS is located at the intergenic region between gIV and the *ori* (Peeters, et al., 1985). Forth, the *Xanthomonas* phage PS sequences exhibit dyad symmetry to form a hairpin structure, but their stem (7-8 bp) are much shorter than Ff or Ike phages (24 and 22 bp, Figure 3F) (Peeters, et al., 1985; Russel and Model, 1989). Fifth, the Ff PS stem has a CGGGT sequence closely followed by a GGTG sequence on the ascending limb (Russel and Model, 1989), while the *Xanthomonas* phage PS stem contain a consensus GGX(A/-)CCG(C/T)G sequence (Figure 3E).

Loss of, or mutations to, ϕLf-UK PS caused a nearly 100 folds decrease of TP reproduction. It raises the question as to how these PS mutants still assembled their TP particles despite the lower efficiencies. It has been shown that TPs of pHV33 (a low copy number plasmid) with the f1 PS was independently packaged from the helper f1 genome. However, pHV33 TPs lacking a functional PS can be encapsidated together with f1, which generated larger virus particles (Russel and Model 1989). This “co-encapsidation” is not uncommon even in the wild type M13 population (2 to 5%) (Beaudoin and Pratt, 1974; Scott and Zinder 1967; Russel and Model 1989). Whether or not co-encapsidation occurs between ϕLf-UK and pORIPS (especially in PS-deficient ones) requires further investigation.

Genetic studies revealed that the residues close to the C-termini of pVII (I27) and pIX (R26 and T30) play a vital role in interacting with the PS (Russel and Model, 1989). Our suppressor screenings also identified that the ϕLf-UK ORF4 N36 mutant can compensate for assembly deficiency of PS-phage, indicating that ϕLf-UK gVII is also involved in phage assembly (manuscript in preparation). Identification of protein factor(s) involved in viral assembly is the next imperative step to understand the evolution and diversity of filamentous phage life cycle.

## Data availability

The complete genome sequences of ϕLf2 and ϕLf-UK phage have been deposited at DDBJ/ENA/GenBank under the accession no. MH218848 and MH206184, respectively.

## Ethical approval

This article does not contain any study with human or animal participants performed by the author.

## Author contributions

**Ting-Yu Yeh**: Conceptualization, Formal analysis, Investigation, Project administration, Methodology, Resources, Supervision; Validation, Visualization; Roles/Writing – original draft **Patrick J. Feehley**: Investigation, Software, Writing – review & editing **Michael C. Feehley**: Investigation, Software, Writing – review & editing, **Vivian Y. Ooi**: Investigation, Writing – review & editing, **Pei-Chen Wu**: Investigation, Software, Visualization **Frederick Hsieh**: Resources, Visualization, **Serena S. Chiu**: Investigation, Validation, **Yung-Ching Su**: Validation, Visualization, **Maxwell S. Lewis:** Investigation, Validation, Visualization, **Gregory P. Contreras**: Validation, Writing – review & editing.

## ACKNOWLEDGMENTS

M.C.F. and P.J.F are scholarship recipients of Olivia Constants Foundation. This work is inspired by Dr. Tsong-Teh Kuo’s (1933-2022) and Yi-Hsiung Tseng’s lifetime achievement of *Xanthomonas* phage study.

## REFERENCES

1. Ahmad AA, Askora A, Kawasaki T, Fujie M, Yamada T. 2014. The filamentous phage XacF1 causes loss of virulence in *Xanthomonas axonopodis pv. citri*, the causative agent of citrus canker disease. Front Microbiol 5:321.

2. Almagro Armenteros JJ, Tsirigos KD, Sønderby CK, Petersen TN, Winther O, Brunak S, von Heijne G, Nielsen H. 2019. SignalP 5.0 improves signal peptide predictions using deep neural networks. Nat Biotechnol. 37:420–423.

3. Bai MS 1989. Studies on insertion of filamentous phage DNA into the host chromosome. Master thesis. National Taiwan University. http://tdr.lib.ntu.edu.tw/jspui/handle/123456789/75692

4. Beaudoin J, Pratt D. 1974. Antiserum inactivation of electrophoretically purified M13 diploid virions: model for the F-specific filamentous bacteriophages J Virol. 13:466–9.

5. Campos J, Martinez E, Suzarte E, Rodriguez BL, Marrero K, Silva Y, Ledon T, del Sol R, Fando R. 2003. VGJϕ, a novel filamentous phage of Vibrio cholerae, integrates into the same chromosomal site as CTXϕ. J Bacteriol 185:5685–5696.

6. Chang KH, Wen FS, Tseng TT, Lin NT, Yang MT, Tseng YH. 1998. Sequence analysis and expression of the filamentous phage ϕLf gene I encoding a 48-kDa protein associated with host cell membrane. Biochem Biophys Res Commun 245:313–318.

7. Conners R, León-Quezada RI, McLaren M, Bennett NJ, Daum B, Rakonjac J, Gold VAM. 2023. Cryo-electron microscopy of the f1 filamentous phage reveals insights into viral infection and assembly. Nat Commun. 14:2724.

8. Dai H, Chiang KS, Kuo TT. 1980. Characterization of a New Filamentous Phage Cf from *Xanthomonas citri*. J. Gen. Virol. 46: 277–289.

9. Dai H, Tsay SH, Kuo TT, Lin YH, Wu WC. 1987. Neolysogenization of *Xanthomonas campestris* pv. *citri* infected with filamentous phage Cf16. Virology 156:313–320.

10. Dai H, Chow TY, Liao HJ, Chen ZY, Chiang KS. 1988. Nucleotide sequences involved in the neolysogenic insertion of filamentous phage Cf16-v1 into the *Xanthomonas campestris* pv. *citri* chromosome. Virology 167:613–620.

11. Day LA, Marzec CJ, Reisberg SA, Casadevall A. 1988. DNA packing in filamentous bacteriophages. Annu Rev Biophys Biophys Chem 17: 509–539.

12. Dotto GP, Enea V, Zinder ND. 1981. Functional analysis of bacteriophage f1 intergenic region. Virology 114:463–73.

13. Endemann H, Model, P. 1995. Location of filamentous phage minor coat proteins in phage and in infected cells. J Mol Biol 250: 496–506.

14. Fu JF, Chang RY, Tseng YH 1992 Construction of stable lactose-utilizing Xanthomonas campestris by chromosomal integration of cloned lac genes using filamentous phage ϕLf DNA. Appl Microbiol and Biotechnol 37:225–229

15. Gonzalez MD, Lichtensteiger CA, Caughlan R, Vimr ER. 2002. Conserved filamentous prophage in Escherichia coli O18:K1:H7 and *Yersinia pestis biovar orientalis*. J Bacteriol 184:6050–6055.

16. Gruber AR, Lorenz R, Bernhart SH, Neuböck R, Hofacker IL. 2008. The Vienna RNA websuite. Nucleic Acids Res. 36(Web Server issue):W70–4.

17. Hallgren J, Tsirigos KD, Pedersen MD, Almagro Armenteros JJ, Marcatili P, Nielsen H, et al. DeepTMHMM predicts alpha and beta transmembrane proteins using deep neural networks. BioRxiv doi: 10.1101/2022.04.08.487609

18. Hay, I.D., Lithgow, T., 2019. Filamentous phages: masters of a microbial sharing economy. EMBO Rep. 20. 10.15252/embr.201847427.

19. Huber KE, Waldor MK. 2002. Filamentous phage integration requires the host recombinases XerC and XerD. Nature 417:656–659.

20. Jia Q, Xiang Y. 2023. Cryo-EM structure of a bacteriophage M13 mini variant. Nat Commun. 14: 5421.

21. Kamiunten H (1995) Integration of filamentous phage Xf2 DNA into chromosomal DNA of *Xanthomonas campestris* pv. *oryzae*. Jpn J Phytopathol 61:127–129

22. Kamiunten H, Wakimoto S. 1980. Effect of the infection with filamentous phage Xf2 on the properties of *Xanthomonas campestris* pv. *oryzae*. Jpn J Phytopathol 47:627–636

23. Koonin EV, Ilyina TV. 1993. Computer-assisted dissection of rolling circle DNA replication. Biosystems. 30:241–68.

24. Kuo TT, Chao YS, Lin YH, Lin BY, Liu LF, Feng TY. 1987a. Integration of the DNA of filamentous bacteriophage Cflt into the chromosomal DNA of its host. J Virol 61:60–65.

25. Kuo TT, Huang TC, Chow TY. 1969 A filamentous bacteriophage from *Xanthomonas oryzae*. Virology 39: 548–555.

26. Kuo, T.T., Lin, Y.H., Huang, C.M., Chang, S.F., Dai, H., Feng, T.Y., 1987b. The lysogenic cycle of the filamentous phage Cflt from *Xanthomonas campestris* pv. *citri*. Virology 156: 305–312.

27. Ledermann R, Strebel S, Kampik C, Fischer HM. 2016. Versatile vectors for efficient mutagenesis of Bradyrhizobium diazoefficiens and Other Alphaproteobacteria Appl Environ Microbiol. 82:2791–2799.

28. Lin, N.T., Chang, R.Y., Lee, S.J., Tseng, Y.H., 2001. Plasmids carrying cloned fragments of RF DNA from the filamentous phage ϕLf can be integrated into the host chromosome via site-specific integration and homologous recombination. Mol. Genet. Genom. 266, 425–435.

29. Lin NT, Tseng YH. 1996. The *ori* of filamentous phage ϕLf is located within the gene encoding the replication initiation protein. Biochem Biophys Res Commun 228:246–251.

30. Lin NT, Wen FS, Tseng YH. 1996. A region of the filamentous phage ϕLf genome that can support autonomous replication and miniphage production. Biochem Biophys Res Commun 218:12–16.

31. Lin NT, You BY, Huang CY, Kuo CW, Wen FS, Yang JS, Tseng YH. 1994. Characterization of two novel filamentous phages of *Xanthomonas*. J Gen Virol 75:2543–2547.

32. Liu J, Liu Q, Shen P, Huang YP, 2012. Isolation and characterization of a novel filamentous phage from *Stenotrophomonas maltophilia*. Arch. Virol. 157, 1643– 1650

33. Liu TJ, Wen FS, Tseng TT, Yang MT, Lin NT, Tseng YH. 1997. Identification of gene VI of filamentous phage ϕLf coding for a 10-kDa minor coat protein. Biochem Biophys Res Commun 239:752–755.

34. Lopez J, Webster RE. 1983. Morphogenesis of filamentous bacteriophage-F1— orientation of extrusion and production of polyphage. Virology 127: 177–193.

35. Mai-Prochnow A, Hui JG, Kjelleberg S, Rakonjac J, McDougald D, Rice SA. 2015. Big things in small packages: the genetics of filamentous phage and effects on fitness of their host. FEMS Microbiol Rev 39:465–487.

36. Peeters BP, Peters RM, Schoenmakers JG, Konings RN. 1985. Nucleotide sequence and genetic organization of the genome of the N-specific filamentous bacteriophage IKe. Comparison with the genome of the F-specific filamentous phages M13, fd and f1 J Mol Biol. 181:27–39.

37. Peng YH. 2002. Studies on the orf155, orf137 and orf102 at the reverse stranded of filamentous phage ϕLf. Master thesis. National Chung Hsing University. https://hdl.handle.net/11296/stcfd2

38. Peng X, Nguyen A, Ghosh D. 2018. Quantification of M13 and T7 bacteriophages by TaqMan and SYBR green qPCR. J Virol Methods 252:100–107.

39. Petrova, M., Shcherbatova, N., Kurakov, A., Mindlin, S., 2014. Genomic characterization and integrative properties of ϕSMA6 and ϕSMA7, two novel filamentous bacteriophages of *Stenotrophomonas maltophilia*. Arch. Virol. 159, 1293–1303.

40. Rambaut A. FigTree, version 1.4.2 [internet]. San Fransisco: GitHub, Inc.; 2021. Available from: https://github.com/rambaut/figtree/releases

41. Reece KS, Phillips GJ. 1995. New plasmids carrying antibiotic-resistance cassettes Gene 165:141–142.

42. Russel, M., Model P. 1989. Genetic analysis of the filamentous bacteriophage packaging signal and of the proteins that interact with it. J Virol 63: 3284–3295.

43. Ryan RP, Vorhölter FJ, Potnis N, Jones JB, Van Sluys MA, Bogdanove AJ, Dow JM. 2011. Pathogenomics of *Xanthomonas*: understanding bacterium-plant interactions. Nat Rev Microbiol. 9:344–55.

44. Solis-Sanchez GA, Quinones-Aguilar, EE, Fraire-Velazquez S, Vega-Arreguin J, Rincon-Enrique, G. Complete Genome Sequence of XaF13, a Novel Bacteriophage of *Xanthomonas vesicatoria* from Mexico. Microbiol Resour Announc. 2020 9:e01371–19.

45. Scott JR, Zinder 1967. Heterozygotes of phage f1. Mol. Biol. Viruses, Proc. Symp. (1966, 1967), pp. 211–218.

46. Shen CK, Ikoku A, Hearst JE. 1979. A specific DNA orientation in the filamentous bacteriophage fd as probed by psoralen crosslinking and electron microscopy. J Mol Biol.127:163–175.

47. Tseng YH, Lo MC, Lin KC, Pan CC, Chang RY. 1990. Characterization of filamentous bacteriophage phiLf from *Xanthomonas campestris* pv. *campestris*. J Gen Virol. 71:1881–4.

48. Vieira, J., Messing, J., 1991. New pUC-derived cloning vectors with different selectable markers and DNA replication origins. Gene 100, 189–194.

49. Waldor MK, Mekalanos JJ. 1996. Lysogenic conversion by a filamentous phage encoding cholera toxin. Science 272: 1910–1914.

50. Wang TW, Tseng YH. 1992. Electrotransformation of *Xanthomonas campestris* by RF DNA of filamentous phage ϕLf. Lett Appl Microbiol. 14:65–8.

51. Webster RE, Grant RA, Hamilton LA. 1981. Orientation of the DNA in the filamentous bacteriophage f1. J Mol Biol. 152:357–74.

52. Wen FS, Tseng YH. 1994. Nucleotide sequence determination, characterization and purification of the single-stranded DNA-binding protein and major coat protein of filamentous phage ϕLf *of Xanthomonas campestris* pv. *campestris*. J Gen Virol 75:15–22.

53. Wen FS, Tseng YH. 1996. Nucleotide sequence of the gene presumably encoding the adsorption protein of filamentous phage ϕLf. Gene. 172:161–162.

54. Williams PH. 1980. Black rot: a continuing threat to world crucifers. Plant Disease. 64: 736–742.

55. Yeh TY. 2017. Complete nucleotide sequence of a new filamentous phage, Xf109, which integrates its genome into the chromosomal DNA of Xanthomonas oryzae. Arch Virol. 162:567–572.

56. Yeh TY. 2020. XerD-dependent integration of a novel filamentous phage Cf2 into the *Xanthomonas citri* genome. Virology 548: 160–167.

57. Yeh TY, Contreras GP. 2021. Viral transmission and evolution dynamics of SARS-CoV-2 in shipboard quarantine. Bull World Health Organ 99:486–495.

58. Yeh TY, Hsieh ZY, Feehley MC, Feehley PJ, Contreras GP, Su YC, Hsieh SL, Lewis DA. 2022. Recombination shapes the 2022 monkeypox (mpox) outbreak. Med. 3:824–826.

59. Yoshizawa S, Kawai G, Watanabe K, Miura K, Hirao I. 1997. GNA trinucleotide loop sequences producing extraordinarily stable DNA minihairpins. Biochemistry 36:4761–7.

