## Supplemental figure and legend for "The packaging signal of *Xanthomonas* integrative filamentous phages"

Supplemental Figure Legend

Figure S1. Characterization of ORF3 and ORF4 of ϕLf-UK and ϕLf2 (ORF43 and ORF38 of ϕLf). (A) Amino acid alignment of ORF4/ORF38 with the gVII of coliphages and *Salmonella* Ike phage. Conserved N-terminal D/E residues, C-terminal R, and amino acids in transmembrane domain (in bold) are labeled in green, red, and purple. (B) Cryo-EM structure of M13 pVII (PDB ID: 8ixl. Jia and Xiang, 2023) and AlphaFold prediction of ϕLf-UK ORF4 structure (UniProt ID: A0A8B4XAY7). Model confidence of AlphaFold is shown in color. (C) Signal peptide prediction of ORF3/ORF43 by SignalP 5.0 (Sec/SPI: standard signal peptides; Sec/SPII: lipoprotein signal peptides; Tat/SPI: Tat signal peptides). The putative lipoprotein signal peptide sequence is labeled in blue. (D) AlphaFold prediction of ϕLf-UK ORF3 protein structure (UniProt ID: A0A0H2X969).

Figure S2. Phylogenetic analysis of amino acid sequence of pVII homologues. Alignments were generated using MAFFT, and the phylogenetic tree was generated in MAFFT using the neighbour-joining method. Bootstrap values for 5 replicates. The phylogenetic tree was visualized using FigTree. Rooting was done by introducing as an outgroup *Stenotrophomonas maltophilia* protein (WP_24079689).

Figure S1

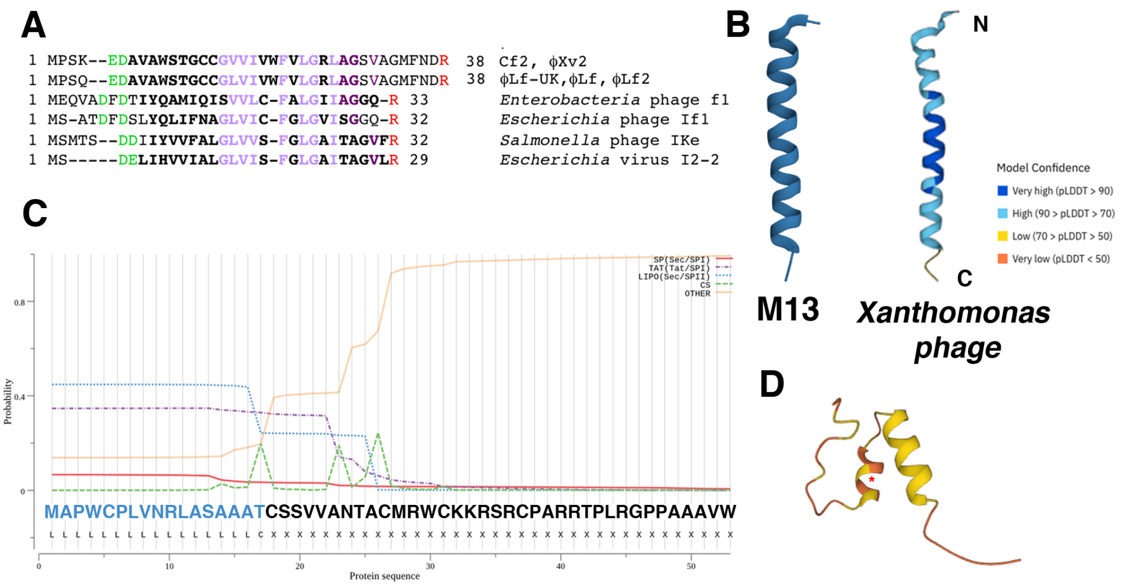
A
1 MPSK--EDAVAWSTGCCGVVIVWFVLGRLAGSVAGMFNDR 38 Cf2, ϕXv2
1 MPSQ--EDAVAWSTGCCGLVIVWFVLGRLAGSVAGMFNDR 38 ϕLf-UK,ϕLf, ϕLf2
1 MEQVADFDTIYQAMIQISVVLC-FALGIIAGGQ-R 33 Enterobacteria phage f1
1 MS-ATDFDSLYQLIFNAGLVIC-FGLGVISGGQ-R 32 Escherichia phage If1
1 MSMTS--DDIIYVVFALGLVVS-FGLGAITAGVFR 32 Salmonella phage IKe
1 MS-----DELIHVVIALGLVIS-FGLGAITAGVLR 29 Escherichia virus I2-2
B
N
C
Model Confidence
Very high (pLDDT > 90)
High (90 > pLDDT > 70)
Low (70 > pLDDT > 50)
Very low (pLDDT < 50)
M13
Xanthomonas phage
C
SP(Sec/SPI)
TAT(Tat/SPI)
LIPO(Sec/SPII)
CS
OTHER
Probability
MAPWCPLVNRLASAAATCSSVVANTACMRWCKKRSRCPARRTPLRGPPAAAVW
Protein sequence
D

Figure S2

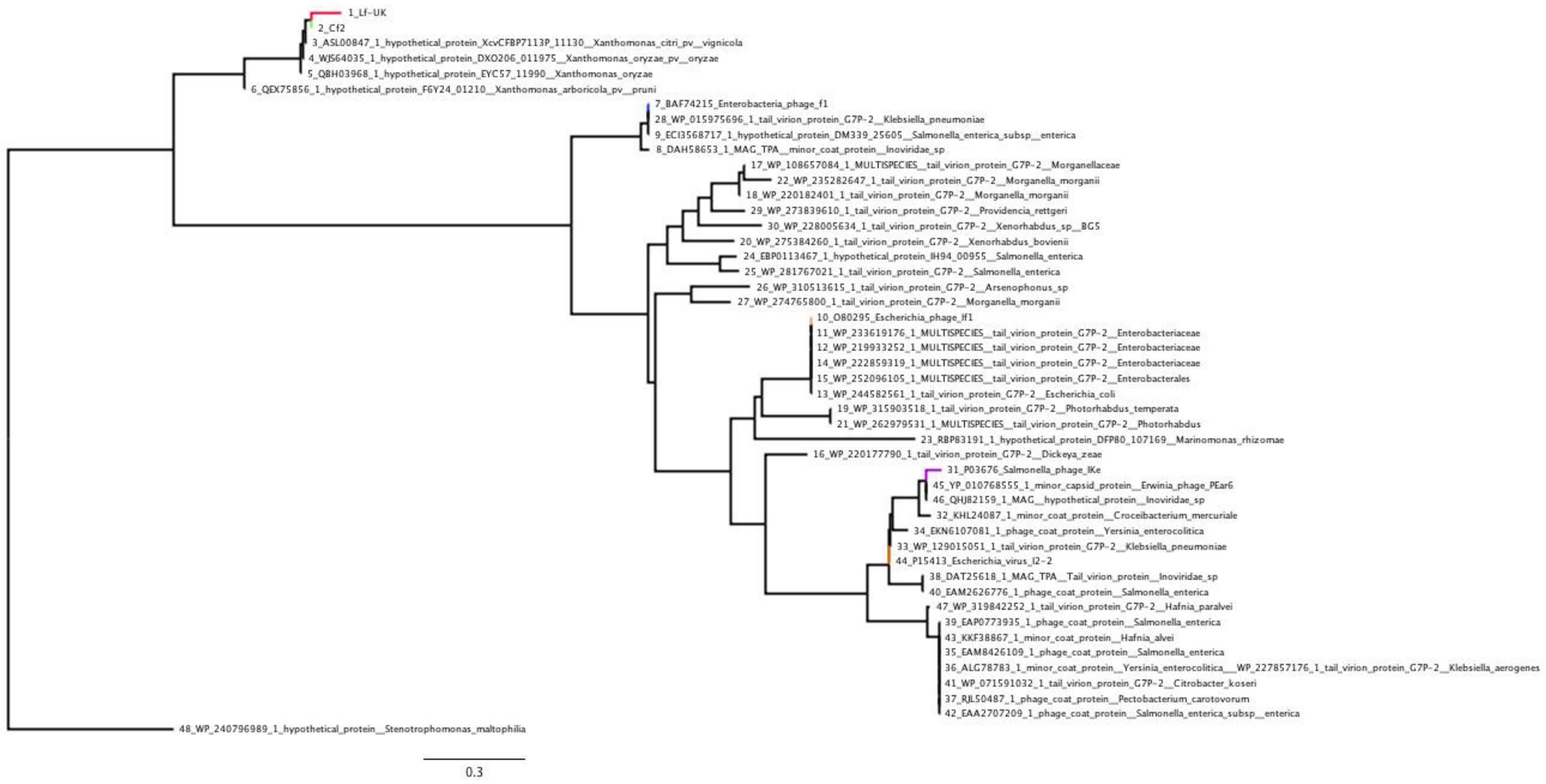
1_Lf-UK
2_Cf2
3_ASL00847_1_hypothetical_protein_XcvCFBP7113P_11130__Xanthomonas_citri_pv__vignicola
4_WJS64035_1_hypothetical_protein_DXO206_011975__Xanthomonas_oryzae_pv__oryzae
5_QBH03968_1_hypothetical_protein_EYC57_11990__Xanthomonas_oryzae
6_QEX75856_1_hypothetical_protein_F6Y24_01210__Xanthomonas_arboricola_pv__pruni
7_BAF74215_Enterobacteria_phage_f1
28_WP_015975696_1_tail_virion_protein_G7P-2__Klebsiella_pneumoniae
9_ECI3568717_1_hypothetical_protein_DM339_25605__Salmonella_enterica_subsp__enterica
8_DAH58653_1_MAG_TPA__minor_coat_protein__Inoviridae_sp
17_WP_108657084_1_MULTISPECIES__tail_virion_protein_G7P-2__Morganellaceae
22_WP_235282647_1_tail_virion_protein_G7P-2__Morganella_morganii
18_WP_220182401_1_tail_virion_protein_G7P-2__Morganella_morganii
29_WP_273839610_1_tail_virion_protein_G7P-2__Providencia_rettgeri
30_WP_228005634_1_tail_virion_protein_G7P-2__Xenorhabdus_sp__BG5
20_WP_275384260_1_tail_virion_protein_G7P-2__Xenorhabdus_bovienii
24_EBP0113467_1_hypothetical_protein_IH94_00955__Salmonella_enterica
25_WP_281767021_1_tail_virion_protein_G7P-2__Salmonella_enterica
26_WP_310513615_1_tail_virion_protein_G7P-2__Arsenophonus_sp
27_WP_274765800_1_tail_virion_protein_G7P-2__Morganella_morganii
10_O80295_Escherichia_phage_If1
11_WP_233619176_1_MULTISPECIES__tail_virion_protein_G7P-2__Enterobacteriaceae
12_WP_219933252_1_MULTISPECIES__tail_virion_protein_G7P-2__Enterobacteriaceae
14_WP_222859319_1_MULTISPECIES__tail_virion_protein_G7P-2__Enterobacteriaceae
15_WP_252096105_1_MULTISPECIES__tail_virion_protein_G7P-2__Enterobacterales
13_WP_244582561_1_tail_virion_protein_G7P-2__Escherichia_coli
19_WP_315903518_1_tail_virion_protein_G7P-2__Photorhabdus_temperata
21_WP_262979531_1_MULTISPECIES__tail_virion_protein_G7P-2__Photorhabdus
23_RBP83191_1_hypothetical_protein_DFP80_107169__Marinomonas_rhizomae
16_WP_220177790_1_tail_virion_protein_G7P-2__Dickeya_zeae
31_P03676_Salmonella_phage_IKe
45_YP_010768555_1_minor_capsid_protein__Erwinia_phage_PEar6
46_QHJ82159_1_MAG__hypothetical_protein__Inoviridae_sp
32_KHL24087_1_minor_coat_protein__Croceibacterium_mercuriale
34_EKN6107081_1_phage_coat_protein__Yersinia_enterocolitica
33_WP_129015051_1_tail_virion_protein_G7P-2__Klebsiella_pneumoniae
44_P15413_Escherichia_virus_I2-2
38_DAT25618_1_MAG_TPA__Tail_virion_protein__Inoviridae_sp
40_EAM2626776_1_phage_coat_protein__Salmonella_enterica
47_WP_319842252_1_tail_virion_protein_G7P-2__Hafnia_paralvei
39_EAP0773935_1_phage_coat_protein__Salmonella_enterica
43_KKF38867_1_minor_coat_protein__Hafnia_alvei
35_EAM8426109_1_phage_coat_protein__Salmonella_enterica
36_ALG78783_1_minor_coat_protein__Yersinia_enterocolitica___WP_227857176_1_tail_virion_protein_G7P-2__Klebsiella_aerogenes
41_WP_071591032_1_tail_virion_protein_G7P-2__Citrobacter_koseri
37_RJL50487_1_phage_coat_protein__Pectobacterium_carotovorum
42_EAA2707209_1_phage_coat_protein__Salmonella_enterica_subsp__enterica
48_WP_240796989_1_hypothetical_protein__Stenotrophomonas_maltophilia
0.3
